# Surface Energy and Viscoelastic Characteristics of *Staphylococcus epidermidis* and *Cutibacterium acnes* Biofilm on Commercial Skin Constructs versus agar

**DOI:** 10.1101/2023.02.10.527933

**Authors:** S. A. Razgaleh, Andrew Wrench, A-Andrew D. Jones

**Affiliations:** Department of Civil & Environmental Engineering, Pratt School of Engineering, Duke University; Duke University Program in Environmental Health; Department of Biomedical Engineering; Thomas Lord Department of Mechanical Engineering & Materials Science, Duke University

**Keywords:** Staphylococcus epidermidis, Cutibacterium acnes, Biofilm, Viscoelastic, Skin Infection

## Abstract

Biofilms are recalcitrant to both study and infectious disease treatment as it requires not only the study or management of single organism behavior, but also many dynamical interactions including but not limited to bacteria-bacteria, bacteria-host, bacteria-nutrients, and bacteria-material across multiple time scales. This study performs comparative and quantitative research of two materials used in biofilm research, TSA agar and skin epidermal, to reveal how adhesion effects viscoelastic properties of biofilms at long time scales. We show that the host surface stressors, such as wettability and surface energy, impact the biofilm’s mechanical integrity and viscoelastic properties. While it is known that the bacteria-material interface influences initial biofilm formation and external stress influences mature biofilm function, this study examines the influence of the bacteria-material interface on mature biofilms. These mechanical viscoelastic properties have the potential to determine metabolite and pathogenesis pathways which means that the platform researchers use to study impacts the outcome.

## I. Introduction

We are headed towards the antibacterial resistance cliff, a point where common medical procedures will run the risk of deadly infections, expected to occur between 2030 – 2050, without much public awareness, funding, or new tools to handle it when failure is estimated to cost over $1 trillion annually.^1, 2^ Of particular concern are acute, chronic infections like diabetic foot ulcers or burn wound infections.^3, 4^ What is promising was a shift in scientific focus from strategies targeting planktonic bacteria to bacterial biofilms.^5-10^ This shift was key because bacteria primarily exist in a biofilm, bacteria grouped together in a viscoelastic matrix of polymers, sugars, proteins, and extracellular DNA, and are more recalcitrant to removal in this state.

Even still, there is less of an understanding of the distinct biofilm phenotypes, relative to planktonic phenotypes and how these phenotypes change over the course of several days. Biofilm phenotypes have been distinguished as a reversible attachment, irreversible attachment, development, maturation, and dispersion.^10^ Attachment occurs when planktonic bacteria approach a surface and adhere to the surface using structures such as pili, fimbriae, or flagella.^11^ Irreversible attachment occurs when the bacterial cytoplasm moves closer to the surface through bacteria cell wall deformation enabling weak attractive Lifshitz-van der Waals forces and when extracellular polymeric substances (EPS) are produced.^12, 13^ Development and maturation have less quantitative distinctions, dependent on levels of EPS, quorum sensing, and other public goods production and levels of metabolic activity. Maturation and dispersion may even be transitory or cyclic complicating this. However, these stages may serve as a better or necessary intervention point. While most interventions and studies of bacteria-material interfaces have focused on the reversible attachment phase, ^14, 15^ here we characterize the mechanical properties as a function of the biofilm-material interface due to the presence of post-attachment biofilms for wound treatment and chronic infections.

Interventions and studies on post-attachment biofilms with respect to skin infections are largely performed on murine or porcine models. Animal models have distinct temperature regulatory systems from humans, different commensal microbiota, and clear bacterial infections faster.^16, 17^ Commercially available skin models are a promising alternative, providing a multilayered immune response for high-throughput screens. Here we compare biofilms grown on a commercially available skin construct to biofilms grown on tryptic soy agar. Tryptic soy agar is designed to be biocompatible, nutrient-rich, hydrated, and hydrophilic. Human skin is a harsh environment for microbes, slightly acidic, oleophilic, with a high neutrophil density. Part of this is due to commensal bacteria organisms. For example, *Staphlyococcus epidermidis* synthesizes phenol-soluble modulins (PSM) using alpha-helical structures, destroying the c1 membrane of pathogens and protecting skin against colonization by foreign bodies.^18^ While *Cutibacterium acnes* produces lipases that hydrolyze the lipids present in sebum and acidify the skin surface by releasing free fatty acids and creating an unfavorable environment for colonization by pathogens. ^18-20^ While most skin bacteria work with immune cells and keratinized skin cells for immune barrier functioning, dysbiosis in the skin microbiome contributes to disease states.^21^

Unlike the attachment phase, most mechanical and viscoelastic measurements of post-attachment biofilms are taken as either dependent functions of bacterial species, nutrient or chemical stress conditions or hydrodynamic conditions, not the material they are grown on. Polymers from biofilms are extracted, purified, and tested for their storage or loss moduli from rheology experiments.^22^ Tensile and compressive strength of those polymers are measured using AFM measurements. Elastic moduli have been measured as functions of overexpression or deficient production of specific EPS components.^23, 24^ Simulations show both the shape (commonly known fluffy or smooth profiles) and removal efficiency are functions of the viscoelastic properties of biofilms,^25, 26^ while we have shown previously that nutrient transport and overall viability is a function of shape and applied shear. ^27^ Furthermore, signaling that modifies biofilm growth and EPS production can also come from both internal cues like quorum sensing compounds and EPS^28-30^and external cues,^31^ including nutrient gradients, hydrodynamic shear gradients,^32^ electrical surface charge,^33^ and nanotexture.

Here, we describe the dependence of viscoelastic properties of post-attachment *Staphylococcus epidermidis* and *Cutibacterium acnes* biofilms on tryptic soy agar and commercially available EpiDerm™ reconstructed human epidermis. We also analyze the storage and loss modulus behavior of deformation in biofilms of these species dependent on surface energy.

## II. Materials & Methods

### a. Skin Culture

Reconstructed human epidermis, Epi-200, were purchased from MatTek. The EpiDerm™ System consists of normal, human-derived epidermal keratinocytes (NHEK) cultured to form a multilayered, highly differentiated model of the human epidermis. Upon receiving the tissues, tissues are transferred into the assay medium via tissue inserts and incubated for 1 h at 37±1°C, 5±1 % CO_2_, and 90% ± 10% relative humidity (RH) in accordance with the manufactures protocol.

**Table 1.** Contact angel measurement and Surface energy calculation

| Substrate Surface | Surface Energy (mJ.m<sup>-2</sup>) | Contact Angle (°) |  |  |
| --- | --- | --- | --- | --- |
|  |  | DI-Water | Ethylene Glyco | 1-Bromonapthalene |
| TSA Agar | 301.23 | 24.94 | 28.04 | 65.04 |
| Skin | 36.5<sup>35</sup> | - | - | - |

### b. Biofilm Cultivation

The commensal bacteria of the skin microbiome, *Staphylococcus. epidermidis* Fussel NCTC 11047, and *Cutibacterium acnes* NCTC 737 (ATCC) are inoculated 24 h before dosing using a single colony at static condition inside a CO_2_ incubator at 27 °C. *S. epidermidis* is cultured in the standard Tryptic soy Broth (Bacto®, BD Difco, US), while *C. acnes* is cultured in Bacto® Thioglycolate broth media (Bacto®, BD Difco, US). Then, substrates (TSA, Epi-200 skin) are dosed with 100 µL of bacteria inoculation. Biofilms are grown statically in 6-well polystyrene tissue culture plates (Avantor) at 24h, 48h, and 72h.

### c. Surface Energy Measurements

Goniometer and sessile drop method were employed to measure the hydrophobicity (wettability) and surface energy of substrates at room temperature. Biofilm was grown on these materials by introducing a 24h inoculated bacteria suspension and 24h incubation at 37±1°C, 5±1 % CO_2_ in the air, and 90% ± 10% relative humidity (RH).

### d. Thickness and Viscoelastic Measurements

Biofilm samples grown on TSA substrate were cut into 8 mm diameter disks. Skin samples were removed from their insert using a sterilized scalpel. Dynamic rheological measurements were carried out on a rheometer (Anton-Paar) using parallel plate PP08/P-PTD200 geometry (8 mm diameter; zero gap) at the room temperature. Measurements were carried out immediately after placing the samples on the plate. All experiments were carried out in three technical and four biological replicates, and slippage of biofilm due to applied stress was carefully avoided by selecting appropriate operating parameters.

## III. Results

### a. Surface Energy & Adhesion

The challenge in targeting biofilms via the extracellular matrix is due to the immense diversity in the mechanisms by which bacteria interact with the material to form biofilms.^34^ Adhesion and wettability (hydrophobicity or hydrophilicity) are directly connected to surface energy. Surface energy reveals information about the interactions of a biofilm with the material at its contact point.

Surface energy analysis of these platforms indicates a significant difference in the surface level on these platforms. Comparing TSA agar with the Epi-200 human epidermis model shows the human epidermis has an order of magnitude lower surface energy. This indicates that surface tension and hence adhesion forces at the bacteria-material interface with human epidermis are lower than the tested TSA platform. Moreover, depending on the hydrophilicity of the bacteria strain and the mechanism used for attachment, the adherence to the surface can significantly vary. For instance, *S. aureus* cells adhere to the hydrophobic surface by many weakly binding macromolecules, while their adherence to the hydrophilic surface is through strong but few macromolecules ^36^. In *S. epidermidis*, the initial attachment is mediated by cell surface proteins that bind to the mammalian extracellular plasma proteins.^13^

The nature of bacteria and the surface interaction dictates their adhesion to the surface and, in turn, the density and distribution of a bacterium colony, especially in the case of multispecies biofilm. Next, we discuss the implication of surface physiochemical properties in biofilm beyond their interfacial interaction using rheometric techniques and investigate their viscoelastic response.

### b. Implication of Viscoelasticity

Once bacteria irreversibly adhere to a material surface, development, maturation, and dispersion all present with different components of EPS. EPS plays an important role in the recalcitrance toward chemical and mechanical changes and in producing phenotypes that differ from their planktonic counterparts ^37^. Mechanical force on the biofilm may exceed the acting force between the different organisms in a biofilm, resulting in an overload and cohesive failure. When the force exceeds the adhesion force between the biofilm and the substratum surface induces adhesive failure, and the biofilm dislodges from the surface. In a biofilm, EPS composition and distribution contribute to its structural integrity and resistance to chemical challenges such as antibacterial and antibiotic treatments. The transport of chemicals and access to nutrients is limited by the composition of EPS. EPS of biofilms is adaptable both in space and time to its environmental conditions, acting as a major impediment to nutrient deprivation resulting in the development of altered phenotypes.^37^ Viscoelastic characteristics of biofilms are a combination of composition and structural properties. Viscoelasticity is a material property that exhibits both viscous and elastic behaviors. Elastic materials deform instantaneously to relieve stress and return to their original state, while viscous materials deform irreversibly over time. Viscoelastic materials deform under stress and return to a similar state to their original state over time, but not identical. Elasticity is the result of bond stretching. On the other hand, viscosity is the result of atomic and molecular flow. Biological materials are complex and cannot be explained by elasticity, viscosity, or a single combination of elasticity and viscosity.

**Figures 2a** and **2b** show the storage and loss modulus, respectively, of *S. epidermidis* and *C. acnes* on TSA. Storage modulus represents the ability of a material to store energy or deformation elastically. While loss modulus represents the ability of a material to dissipate stress as a heat. Additionally, the storage modulus (G’) and loss modulus (G”) describes the rigidity and fluidity of the material.^38^The viscoelastic behavior of biofilm and its impact on the integrity of the growth platform is dependent on the bacteria species and unique to its interaction with the substrate. This is likely due to the composition of each biofilms’ EPS composition. Both *S. epidermidis* and C. acnes biofilm showed a reduction of the storage and loss modulus when grown on TSA. However, there was a greater reduction of storage and loss modulus for *S. epidermidis* than the *C. acnes* biofilm on TSA. On the other hand, as shown in Figure 3a and 3b, *S. epidermidis* growth on Epi-200 reduced the storage and increased the loss modulus while *C. acnes* growth reduced both the storage and loss moduli compared to the Epi-200.

**Figure 1a.**
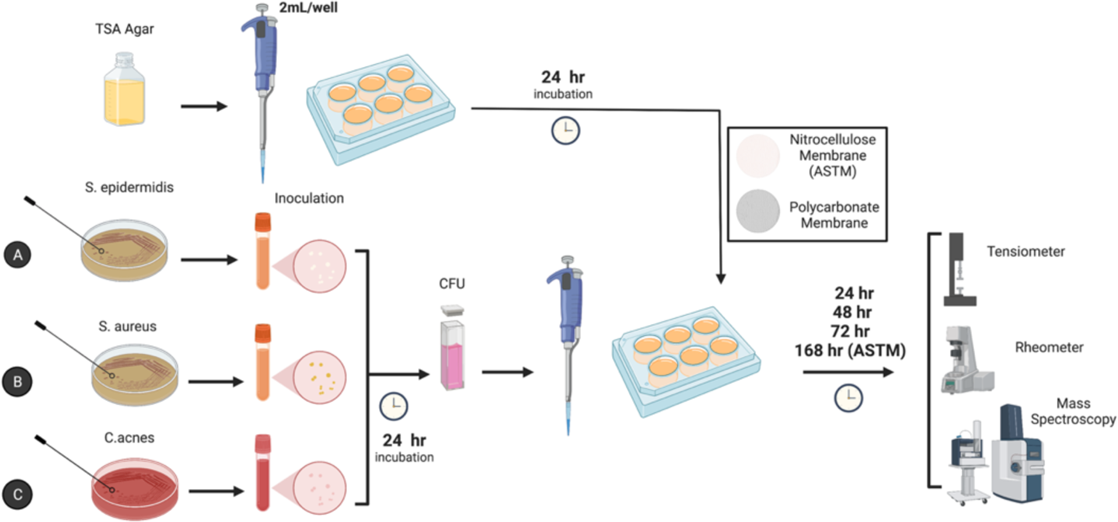
Experimental methodology for measuring physical properties of bacteria biofilms.

**Figure 1b.**
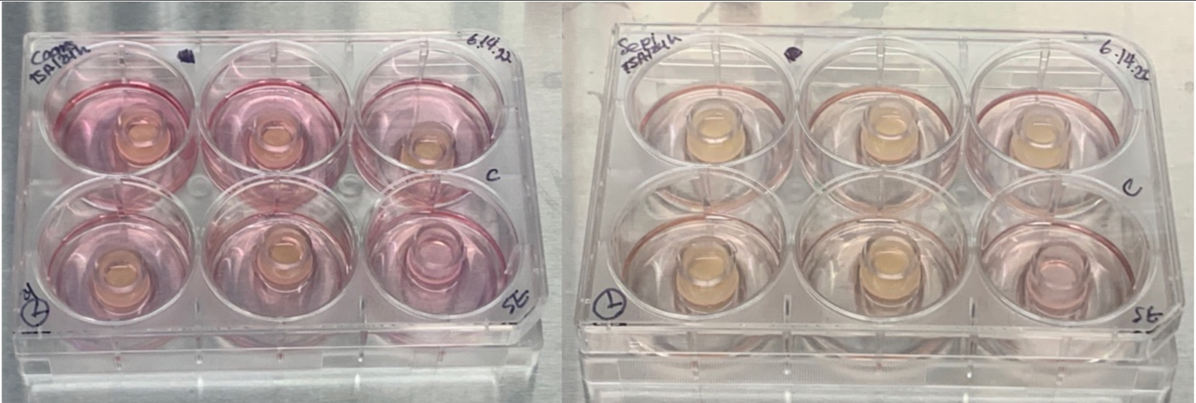
*C. acnes* and *S. epidermidis* on EpiDerm™ human epidermis in 6-well polystyrene culture plates.

**Figure 2.**
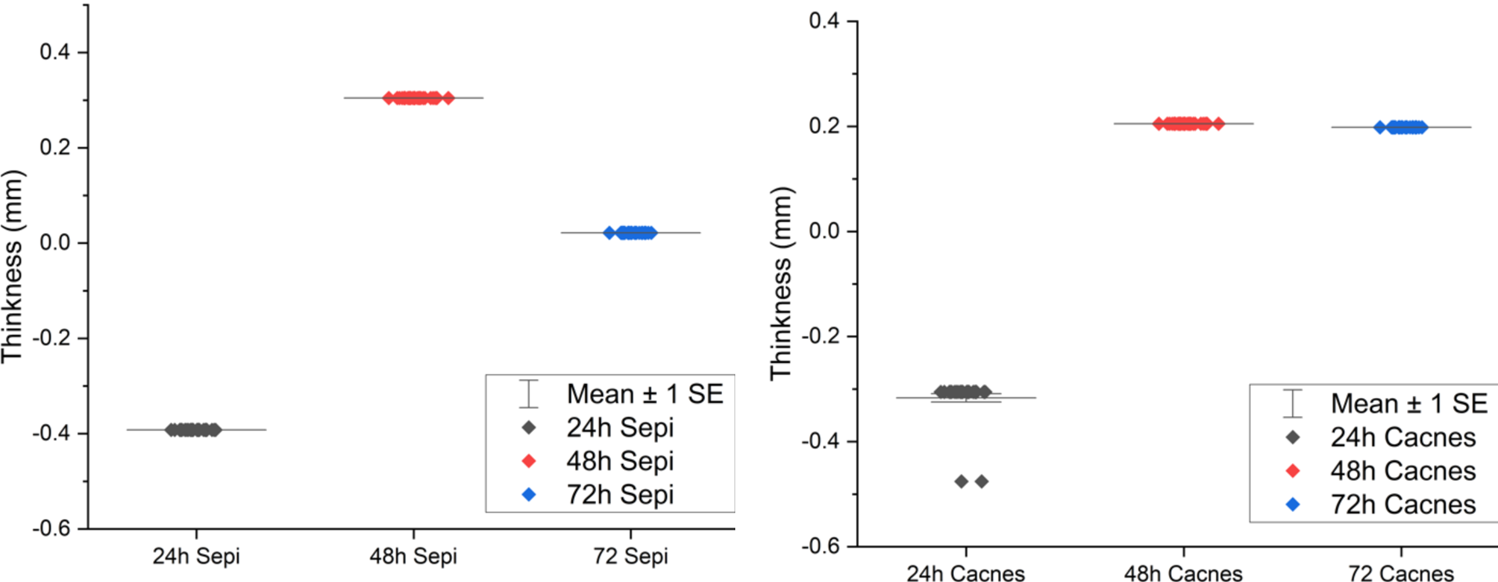
Thickness of biofilms for *S. epidermidis* and *C. acnes* on TSA made using a rheometer. Results reported relative to level surface of the same uncultured agar.

**Figure 2a.**
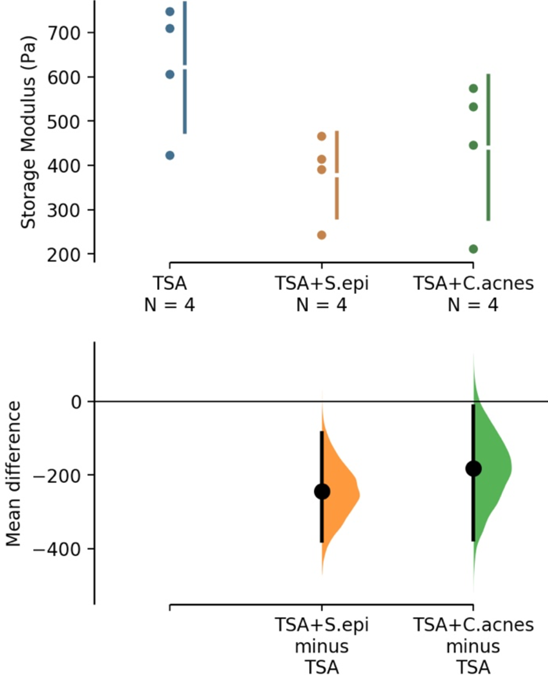
Storage modulus of *S. epidermidis* and *C. acnes* grown on TSA. Each data point is a biological replicate (*n* = 4) average of *n* = 3 technical replicates.

**Figure 2b.**
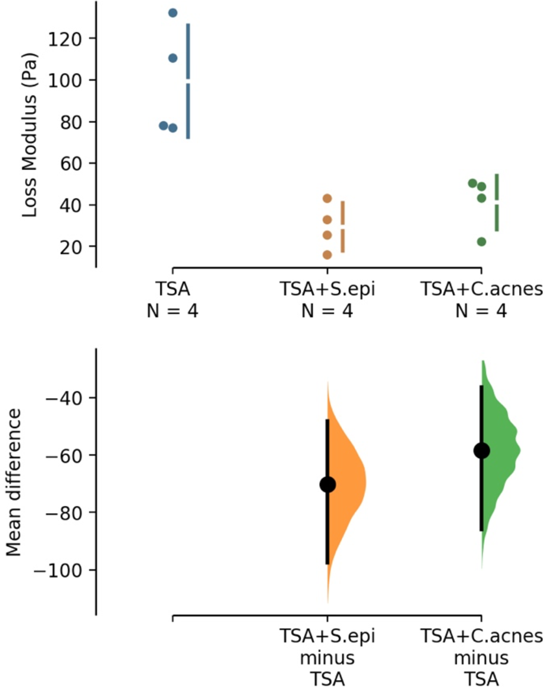
Loss modulus of *S. epidermidis* and *C. acnes* grown on TSA. Each data point is a biological replicate (n = 4) average of *n* = 3 technical replicates.

**Figure 3a.**
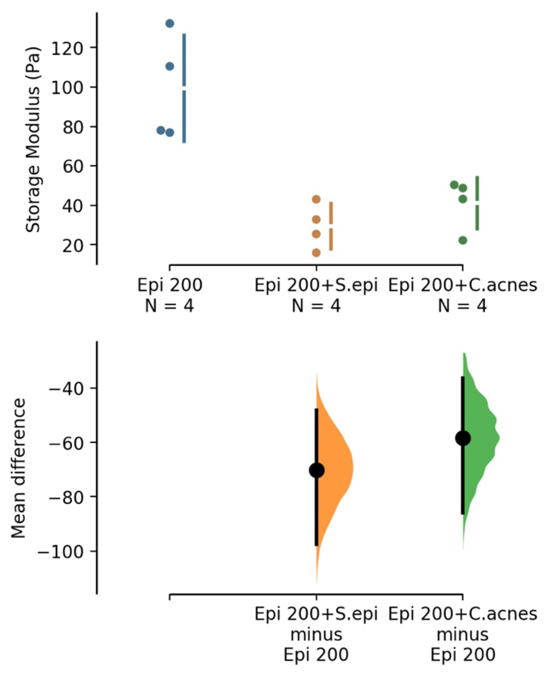
Storage modulus of *S. epidermidis* and *C. acnes* grown on Epi-200. Each data point is a biological replicate (***n* = 4**) average of (***n* = 3**) technical replicates.

**Figure 3b.**
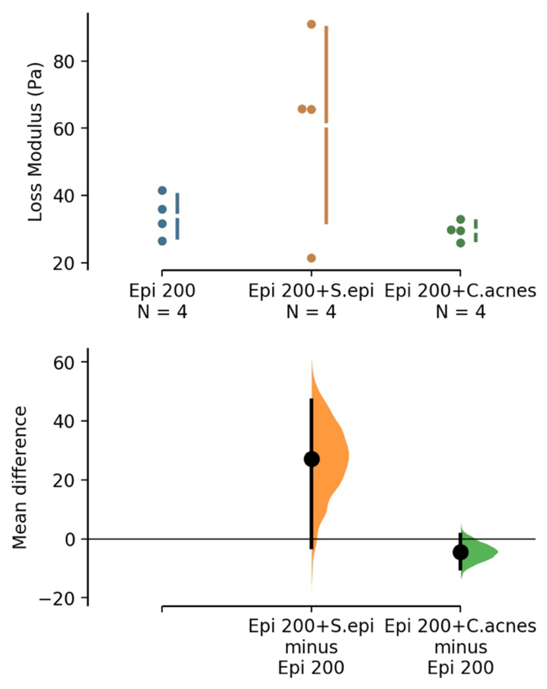
Loss modulus of *S. epidermidis* and *C. acnes* grown on Epi-200. Each data point is a biological replicate (***n* = 4**) average of (***n* = 3**) technical replicates.

## IV. Discussions

Biofilm development generally follows a process of reversible adhesion, irreversible adhesion, development maintenance, and dispersion to form biofilm.^10, 39^ Most research on bacteria-material interaction focuses on adhesion, reversible or irreversible.^37, 40-50^ Experimental timepoints in this experiment are targeted towards the development and maintenance stages of biofilm growth, what we collectively refer to as post-attachment, in interaction with specific materials. The comparison of rheological behavior of biotic and abiotic materials in these studies are aimed at providing indicators of specific material-biofilm interactions. Rheological behavior, represented by storage and loss moduli are aimed at ultimately representing the viscoelasticity of biofilms.^38^

Storage and loss moduli are part of bacterial biofilms’ viscoelastic properties, and thus their capability to resist mechanical deformation. Ultimately, the viscoelastic properties of a biofilm, are a measure of the biofilm’s capacity to demonstrate elastic behavior, and revert to its shape before a force application, or flow with viscosity according to shear stress application. While heterogenous micro-environments within a biofilm complicate the understanding of viscoelastic measurements and generalizations for a biofilm, viscoelasticity is a strong indicator for mass transport models within these regions of a biofilm. Demonstrated in our experiments, unique material surface interactions lead to mechanically different biofilms, indicating transport of infectious disease treatment conditions, and model systems aimed at studying infectious disease treatment, may vary significantly in accuracy depending on material surface structure, especially structures of biotic versus abiotic materials. Research in the field of biofilm adhesion has proven differential growth patterns of biofilms dependent on nanostructure and roughness, or curvature of a surface.^51^ Distinct growth patterns of our bacterium, in particular *C. acnes* in response to the implementation of several variables, including the morphology of keratinocytes and fibroblasts in culture according to EpiDerm™ tissue indicates skin commensal strains within our experiments may also vary in growth depending on these factors.

Mass transport models of biofilms, typically understood as a quantification of elastic behavior, which is then associated diffusion through a solid, versus that of viscous flow through a fluid,^52^ a biofilm’s primary models for mass transfer, are essential for studies in the movement of treatment antibiotics and nanoparticles, immune cells, proteins or nutrients. The implementation, however, of analyzing a biofilm with respect to rheology, along with imaging of skin commensals in culture with keratinocytes and fibroblasts, would be necessary to provide novel insight on the mechanical properties of bacteria that establish commensal relationships with the human epidermis.

Particularly, *Cutibacterium acnes* demonstrably minimizes variation in viscoelastic behavior when compared to viscoelastic behavior of its other skin commensal counterpart, *Staphylococcus epidermidis*, and is provided with a platform for biofilm development in the presence of specific attachment with commercially available skin cell models. Given *C. acnes* and its presence as an anaerobe often associated with opportunistic pathogen behavior^53^ we observe that *C. acnes* specific maintenance with skin is an essential component of its ability to form biofilms, where biofilm behavior and cell proliferation is most predictable. Given *C. acnes’* fledgling capacity within our experimental parameters to develop biofilms and proliferate at all in anaerobic conditions without the implementation of mammalian cells, and the maintained growth of *C. acnes* in planktonic cultures, it can reasonably be inferred that there is an essential component of *C. acnes* biofilm development represented in mammalian cells cultures of EpiDerm™. While the mechanism by which *C. acnes* proliferates in the presence of human epidermal keratinocytes and fibroblasts may also be an element represented in other systems for attachment, this carries great implications in both the fields of basic microbiology, as well as medical health applications. Reduction of *C. acnes* biomass may lie in modification of surfaces, such as those involved in medical device implantation that do not conserve those elements of mammalian keratinocytes and fibroblasts in culture that provide adequate surfaces for biofilm development. Further analyses on whether *C. acnes* demonstrates these patterns on keratinocyte and fibroblast surfaces that do not mimic surface morphology of *in vivo* experiments, as EpiDerm™ does, and comparison of conserved transcriptomic and proteomic signatures of *C. acnes* cells in biofilms would allow research to further support the conclusion and distinctions on whether biofilm formation for this bacterium is predominantly as a function of mechanical behavior and attachment surface potential, or rather the actual production of proteins in response to metabolites in mammalian cells, or some other variable. Given the commonality of *C. acnes* and *S. epidermidis* in the commensal microbiota, there exists extensive genetic classification, and so analysis of the transcriptome would be likely to reveal variation in well-studied proteins, whose functions are understood. Development of biofilms in similar commensals often results in stark differences related to metabolism, efflux pumps, and other genes related to adhesins. The viscoelastic behavior here, of low standard error also demonstrate that *C. acnes* more than likely demonstrates comparatively more homogenous biofilm structures and microenvironments, than, say, *S. epidermidis*, whose variation may reasonably be due to regions of varying viscoelasticity, which our methods would not have robustly identified.

While definite trends in biofilm development on different surfaces, particularly in *C. acnes* cultures which do not form significant biofilm in our experimental conditions outside of commercially available skin, show that biofilms do in fact respond to their environment, our hypotheses are broad and awaiting further experimentation that may provide a key element to further understanding biofilm antibiotic resistance within biofilms. Furthermore, explicit distinction between biofilm growth patterns and genetic signaling allow us to more robustly define biofilm-similar genetic profiles for the identification of biofilms with phenotypes that carry an increased likelihood to slow metabolism, conduct gene transfer, and up-regulate efflux pumps prior to accumulation of biofilm-associated metabolites, which is especially important in those contexts within pores or wound beds where the accumulation of these metabolites and specific EPS structures may not be feasible.

Biofilm formation in chronic and acute wounds has made medical treatment very challenging due to disruption in the innate immune system and the penetration body’s physical barrier. Most common treatments are ineffective in eradicating biofilm or preventing its formation, implying the need for innovative and effective techniques. Recently, the nanotechnology-based mechanism for drug delivery has been the focus of treatment for wound and implant biofilm infections^11^. Moreover, bacteria can exhibit a symbiotic relationship with their hosts and are in constant communication with the host immune system, hence involved in wound healing ^21^. However, other studies have suggested contradictory data; for instance, the absence of commensal skin organisms has improved macroscopic wound healing and closure rate ^21, 54^.

While the results presented in this paper cannot approximate the implications of biofilm on the skin for wound healing applications, it does show the importance of more skin like models for representing properties of biofilm infections. Future skin models should consist of multiple highly regulated but interdependent phases of inflammation, proliferation, epithelialization, angiogenesis, remodeling, and scarring ^11, 55^ Then studies of bacterial biofilm infections can be integrated with disruption during this cascade and more proper methods of healing and treatment can be pursued.^11^

## V. Conclusion

We showed that the bacteria-material interface impacts the viscoelastic characteristic of biofilms post-attachment. We found that the surface energy of materials commonly used for biofilm research vary significantly from that of commercially available reconstructed human epidermis, an order of magnitude less. However, while the magnitude difference in storage modulus for *S. epidermidis* is consistent with the drop in surface energy across materials, the variation in loss modulus is not consistent. Furthermore, the variation in storage and loss modulus for *C. acnes* properties is not consistent across material platforms.

## Acknowledgments

Research reported in this publication was supported by the National Institute of General Medical Sciences of the National Institutes of Health under Award Number NIH R35 GM142898 and by the National Institute of Environmental Health Sciences of the National Institutes of Health under Award Number T32ES021432 (Duke University Program in Environmental Health). The content is solely the responsibility of the authors and does not necessarily represent the official views of the National Institutes of Health.

## Notes

### Competing Interest Statement

The authors have declared no competing interest.

### Summary of Updates

Updated ORCID and name formatting for the second author.

